# Application of long-interval paired-pulse transcranial magnetic stimulation to motion-sensitive visual cortex does not lead to changes in motion perception

**DOI:** 10.1101/766428

**Authors:** Olga Lucia Gamboa Arana, Alexandra Brito, Zachary Abzug, Tracy D’Arbeloff, Lysianne Beynel, Erik A. Wing, Moritz Dannhauer, Hannah Palmer, Susan A. Hilbig, Courtney A. Crowell, Rachel Donaldson, Roberto Cabeza, Simon W. Davis, Angel V. Peterchev, Marc A. Sommer, Lawrence G. Appelbaum

**Affiliations:** Department of Psychiatry and Behavioral Sciences, Duke University School of Medicine; Department of Biomedical Engineering, Duke University; Department of Psychology & Neuroscience, Duke University; Department of Neurology, Duke University School of Medicine; Department of Electrical & Computer Engineering, Duke University; Department of Neurosurgery, Duke University School of Medicine; Department of Neurobiology, Duke University School of Medicine

**Author notes:** **Corresponding author:** L. G. Appelbaum, 400 Trent Dr., Durham, NC, 27710.

**Keywords:** Transcranial Magnetic Stimulation, Visual Motion, Paired Pulse TMS, Motion Sensitive Cortex, hMT+

## Abstract

The perception of visual motion is dependent on a set of occipitotemporal regions which are readily accessible to neuromodulation. Previous studies using paired-pulse Transcranial Magnetic Stimulation (ppTMS) have provided evidence of the capacity of this type of protocols to modulate cognitive processes. To test whether such cortical modulation can be observed in the visual system, particularly during motion perception, ppTMS was applied to the occipital cortex using both scalp-based and meta-analytic targeting coordinates. In this within-subject, sham-controlled study, fifteen subjects completed two sessions in two consecutive weeks. On the first visit, subject-specific resting motor threshold (RMT) was determined and participants performed an adaptive motion discrimination task to determine individual motion sensitivity. During the second visit, subjects performed the same task with three individualized difficulty levels as two TMS pulses were delivered respectively −150 and −50 ms prior to motion stimulus onset at 120% RMT, under the logic that the cumulative inhibitory effect of these two pulses would alter motion sensitivity as measured by the individually calibrated task. The ppTMS was delivered at one of two locations: 3 cm dorsal and 5 cm lateral to inion (scalp-based coordinate), or at the site of peak activation for “motion” according to the NeuroSynth fMRI database (meta-analytic coordinate). Sham stimulation was delivered on one-third of trials and evenly between the two targets. Analyses showed no significant active-versus-sham effects of ppTMS when stimulation was delivered to the meta-analytic (p = 0.15) or scalp-based coordinates (p = 0.17), which were separated by 29 mm on average. Additionally, there was no was significant interaction between ppTMS at either location and task difficulty level (p = 0.12 and p = 0.33, respectively). These findings fail to support the hypothesis that long-interval ppTMS recruits inhibitory processes in motion-sensitive cortex, but must be considered within the limits of the current design choices.

**HIGHLIGHTS:**

- Long-interval paired-pulse TMS was applied to visual cortex during a motion task
- The ppTMS was delivered according to scalp and meta-analytic coordinates, as well as sham
- No effects of active-versus-sham stimulation were observed on motion task performance

## INTRODUCTION

The visual system provides an attractive target to investigate the capacity of non-invasive neuromodulation to affect cortical representations. Going beyond its more typical use in studying motor evoked potentials (MEPs) in hand muscles, transcranial magnetic stimulation (TMS) that targets visual perception can take advantage of the wide array of possible representations and processes undertaken by the visual system. One quality of the visual system which has been explored extensively is the capacity to perceive motion. In particular, area hMT+, the human homologue to macaque medial temporal cortex, is a relatively superficial cortical region that contains a robust representation of stimulus motion [1, 2].

While the neuromodulation of a number of visual functions have been extensively mapped, targeting motion-sensitive cortex remains challenging, with highly-variable rates of behavioral engagement as measured with phosphene or motion-phosphene (‘mophene’) induction [3, 4]. As such, it is unclear what approach is best to properly target this function in human subjects. Here we investigated this question through application of paired-pulse transcranial magnetic stimulation (ppTMS) to motion-sensitive visual cortex. Using two different ppTMS targeting approaches, we applied the neurostimulation just before subjects had to make individually calibrated motion coherence judgments. The overall goal was to determine if the ppTMS disrupted those judgments.

Neuromodulatory effects of ppTMS have been extensively demonstrated in the motor cortex [5–11] as well as in other non-motor cortical areas. For example, ppTMS to the parietal cortex has been reported to modulate excitability and behavior during tactile and visuospatial perception tasks [12, 13] while stimulation to the dorsolateral prefrontal cortex (DLPFC) during encoding modulates retrieval and working memory processes [14–16]. Within the domain of vision, ppTMS over the primary visual cortex has been reported to improve phosphene perception [17], modulate perceptual decision making [18], and disrupt motion perception [19–21] and motion prediction [22].

The characteristics of these observed effects are highly dependent on temporal and spatial parameters defining the ppTMS protocol. For example, the prior studies used stimulus onset asynchronies (SOAs) between the ppTMS and the onset of visual motion that ranged from −42 ms [20] to 100 ms [23], typically attempting to modulate either baseline excitability when the visual stimulus arrives or alter a specific process in the neuronal cascade. Similarly, the inter stimulus interval (ISI) between pulses plays an important role, with past literature in the motor cortex showing inhibitory effects with ISIs between 50 and 200 ms [24], maximal inhibition at 100 ms ISI [25], and previous ppTMS applications outside the motor cortex adopting ISIs within this range [18, 25, 26]. Moreover, studies of this kind relied on a variety of targeting techniques, including scalp landmarks (e.g., inion), functional mapping based on fMRI or EEG data (i.e., “functional localizers”) [21, 27, 28], and identification of phosphenes and/or mophenes.

Extending this past literature, the current study applied long-interval (100 ms) ppTMS to the left hMT+ immediately before a near-threshold motion discrimination task presented to the right visual field. It was hypothesized that ppTMS would cause behavioral impairment in motion perception, driven by accumulated inhibitory effects of the paired pulses that were delivered before the motion stimulus presentation. In particular, we hypothesized that the task would be impacted by long-interval cortical inhibition effects, which occur 50–200 ms after suprathreshold TMS pulses [5, 29]. Moreover, greater modulatory effects were expected for meta-analytic targeting relative to the scalp-based targeting approach, as it scales with head size. To test this, 15 healthy young adult subjects completed a motion discrimination protocol during which active or sham ppTMS, with an ISI of 100 ms, was applied 50 ms before onset of the visual stimuli to motion-sensitive cortex using two different targeting approaches: a traditional 3-cm by 5-cm scalp-based approach using scalp landmarks and a functional approach using meta-analytic information from fMRI studies of motion perception that scale approximately to the size of the participants head. As such, this study aimed to test ppTMS on a calibrated motion sensitivity task to determine if modulatory effects could be observed.

## MATERIAL AND METHODS

### Participants

Twenty-two healthy young adults were recruited and provided written informed consent for the present study, which received approval by the local University Institutional Review Board (Pro00082433) and was pre-registered on ClinicalTrials.gov (NCT03259568). Of the 22 participants who provided consent, four were excluded due to poor performance on the behavioral task, two for contra-indications to TMS, and one who voluntarily withdrew from the protocol. Therefore, data from 15 participants (8 females) who completed the full protocol was used for these analyses. These participants had a mean age of 22.7 years (SD = 3.8) and had normal or corrected-to-normal vision. Participants reported no neurological or psychological disorders and were all native English speakers. Participants were compensated $20 per hour for their time.

### Protocol

Participants enrolled in a two-visit protocol, with an average of 3.2 (SD = 2.9) days between visits. During the first visit, participants responded to a TMS safety questionnaire and underwent a urine drug and pregnancy screening to determine eligibility. Resting motor threshold (RMT) was then determined, after which participants were trained on the motion discrimination task for which they completed between 660 and 1000 trials to reach asymptotic performance. No stimulation was administered during these practice blocks; however, the device was sending pulses in the air at a distance of several feet from the participant to acclimate them to the clicking sounds produced by the coil. On the second visit, after completing one block of practice without stimulation, participants then performed the task at three difficulty levels that were determined on the basis of their individual performance during the first session. During these trials, participants received online paired-pulse TMS at two locations, randomly interleaved with a sham condition. For the sham condition, the coil was flipped such that the edge of the coil was perpendicular to the head, so that a significantly lower magnetic field entered the brain while still inducing a similar auditory sensation. These procedures are described in greater detail below.

### Motion discrimination task

Each trial of the motion discrimination task (**Figure 1**) began with presentation of a fixation cross on a computer monitor. The subjects were required to foveate the cross and then, 750 to 1500 ms after fixation cross appearance, moving dots appeared in the right periphery. The center of the dot field was located 8° from the fixation cross and extended 12° in diameter. During the motion interval, dots moved either upwards to the right or upwards to the left at a 45° angle and remained on the screen for 150 ms. Difficulty was manipulated by altering the coherence of the dot motion (i.e. the percentage of dots moving in the target direction). On the first visit, dot coherence was systematically altered according to a staircase schedule. At the beginning of each block, coherence would start randomly at 0 or 100 percent coherence. After correct decisions, the coherence decreased by 0.2 times the difference between the coherence on that trial and 0%. After errors, the coherence increased by 0.1 times the difference between 100% and the coherence on that trial. Participants were asked to respond as quickly and accurately as possible. Feedback was given after each trial, indicated with a green or red fixation cross for correct or incorrect responses, respectively. The task was identical during the second visit, except that instead of an adaptive staircase procedure, three fixed coherence levels were presented that corresponded to 60% (hard), 75% (medium), and 90% (easy) accuracy according to individual performance during the first visit.

**Figure 1.**
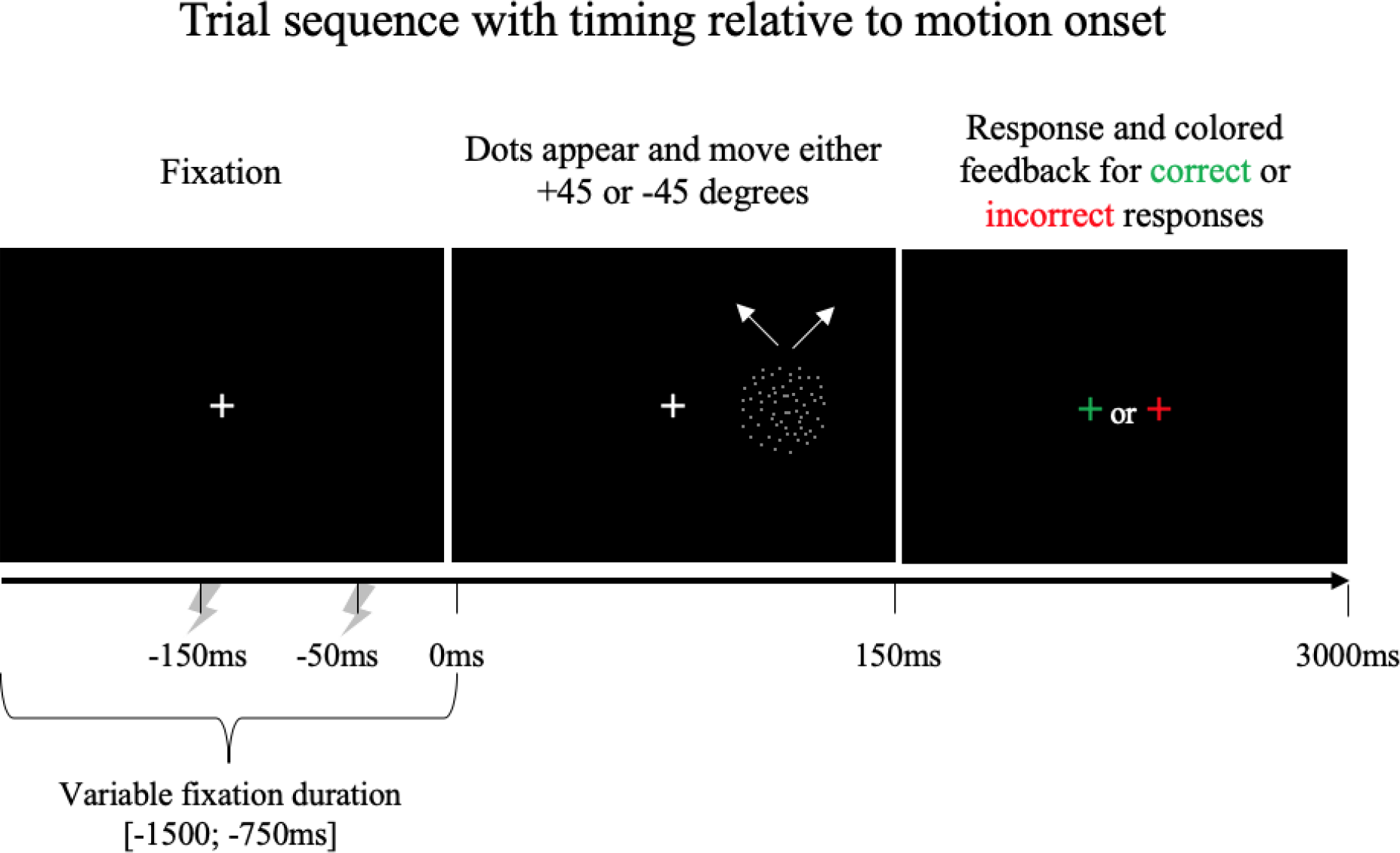
Schematic of trial sequence. On each trial a central fixation was presented, followed by dots in the right visual field that moved either diagonally to the upper left or upper right for 150 ms. Participants then had 3 seconds to respond, and received feedback based on a change of the color of the fixation cross. During visit 1, no TMS was applied. During visit 2, ppTMS was applied with pulses occurring 150 ms and 50 ms prior to motion onset.

### TMS Procedures

All TMS procedures used a Cool-B65 butterfly (figure 8) coil and a MagPro R30 stimulator (MagVenture, Denmark) guided by a Brainsight stereotaxic neuronavigation system (Rogue Research, Canada).

#### Resting Motor Threshold

Resting motor threshold (RMT) was determined for each participant during the first visit. During this procedure, the hot spot was determined for left motor cortex that optimally elicited an MEP in the right first dorsal interosseous (FDI) muscle. The resting motor threshold was then assessed as the lowest TMS pulse intensity that produced, on average, an MEP of 50 μV peak-to-peak amplitude according to a maximum likelihood estimator (TMS Motor Threshold Assessment Tool, MTAT 2.0, http://www.clinicalresearcher.org/software.html)

#### Paired-Pulse Stimulation

Online ppTMS was administered at 120% of RMT over motion-sensitive cortex. The first TMS pulse was delivered 150 ms prior to visual presentation of moving dots, followed by an inter-pulse interval of 100 ms. Therefore, the second TMS pulse occurred 50 ms before onset of the visual motion stimulus. Thus, the collective effects of the two pulses were expected to approximately span the interval before and during the motion discrimination task. As noted above, to control for nonspecific effects due to the clicking sound on task performance, sham stimulation was delivered by tilting the coil 90° perpendicular to the surface of the head while all other parameters described for the active conditions were held constant.

#### TMS Targeting

ppTMS was delivered to the left hemisphere of the motion-sensitive cortex, as this has been shown to produce visual-perceptual effects more consistently than stimulation of the right hemisphere [21]. Two spatial targeting approaches were used. First, in line with numerous studies targeting visual motion cortex in humans, a scalp measurement (the ‘3-5 cm’) procedure was used [27, 28, 30]. After a manual search for the inion, a marker was placed using neuronavigation three centimeters dorsal and five centimeters lateral to the inion. This measurement was marked in the same fashion for all participants and did not take into account individual differences in head size and shape.

A second method of targeting was derived from coactivation information derived from the meta-analytic platform Neurosynth (www.neurosynth.org). Neurosynth is a web-based, publicly accessible database of blood-oxygen-level dependent (BOLD) activations created from thousands of prior fMRI studies [31]. To derive a target from this database, a query was made to Neurosynth using the term “motion” which produced a meta-analysis synthesis of 383 studies, returning a forward probability map in standardized Montreal Neurological Institute (MNI) space. This map was subsequently scaled to each participant’s head size by registering the left and right preauricular points and nasion in Brainsight and warping the Neurosynth map to the standard head model within the Brainsight software. This map was applied as an overlay on the standardized brain and the activation threshold increased to ascertain the peak activation, which was then defined as the second target. The different TMS conditions (scalp-based or meta-analytic TMS coordinates, and sham) were delivered in a randomized order.

### Data Analysis

All statistical analyses were performed in MATLAB (MathWorks, Natick, MA, USA). We analyzed trial-by-trial accuracy (correct = 1, incorrect = 0) and reaction time using logistic mixed effects regression and linear mixed effects regression, respectively. Since motion coherences were selected to normalize accuracy across subjects, we treated difficulty as a continuous fixed effect independent variable (hard = 1, medium = 2, easy = 3). TMS location was treated as a categorical fixed effect independent variable. In addition to main effects, both models featured difficulty × TMS location interaction terms. Participant ID was used as a random effect in both models. We used a post-hoc Bonferroni correction to correct for the n = 2 statistical models used, leading to a significance threshold of p = 0.025.

## RESULTS

### TMS Effects

In general, TMS did not show an effect on behavior at either brain location during the motion discrimination task. Analyses performed using a logistic mixed effects regression demonstrated that TMS did not affect motion discrimination accuracy at the Neurosynth location (main effect: β = 0.189, p = 0.15; interaction with difficulty: β = −0.185, p = 0.12) or at the 3-5 location (main effect: β = −0.182, p = 0.17; interaction with difficulty: β = 0.121, p = 0.33). TMS also did not affect reaction time at the Neurosynth location (main effect: β = 0.020, p = 0.22; interaction with difficulty: β = −0.012, p = 0.28) or at the 3-5 location (main effect: β = 0.013, p = 0.39; interaction with difficulty: β = 0.004, p = 0.729). Significant main effects of difficulty were present for both accuracy (β = 0.986, p < 0.0001) and reaction time (β = −0.056, p < 0.0001) with means and standard deviations shown in **Figure 2**.

**Figure 2.**
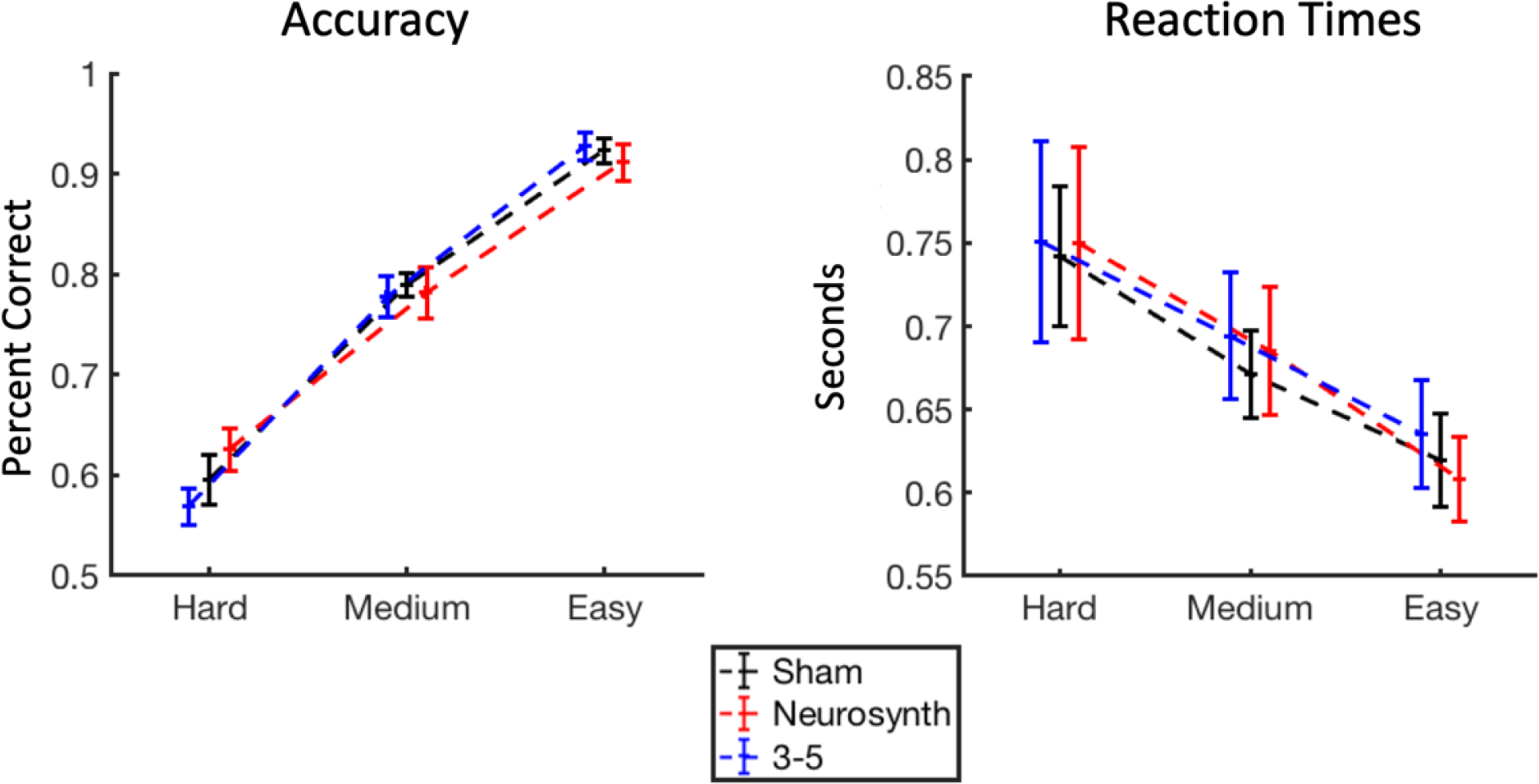
Behavioral performance. Mean accuracy and reaction times, and standard error bars for sham stimulation (black), Neurosynth targeting (red) and 3-5 targeting (blue), shown across difficulty levels.

### TMS Targeting

Cortical targeting of TMS has improved greatly to create more accurate localization of desired neuromodulation. While neither of the approaches used in this study represent the state-of-the-art (e.g. fMRI-localized with electric-field modeling [32]), they do offer a contrast of between a common method that does not scale with head size (3-5 approach) and one that does (Neurosynth approach). As shown in **Figure 3**, variation between the methods was sizable, with a mean distance difference of 29 mm and a spread of several centimeters.

**Figure 3.**
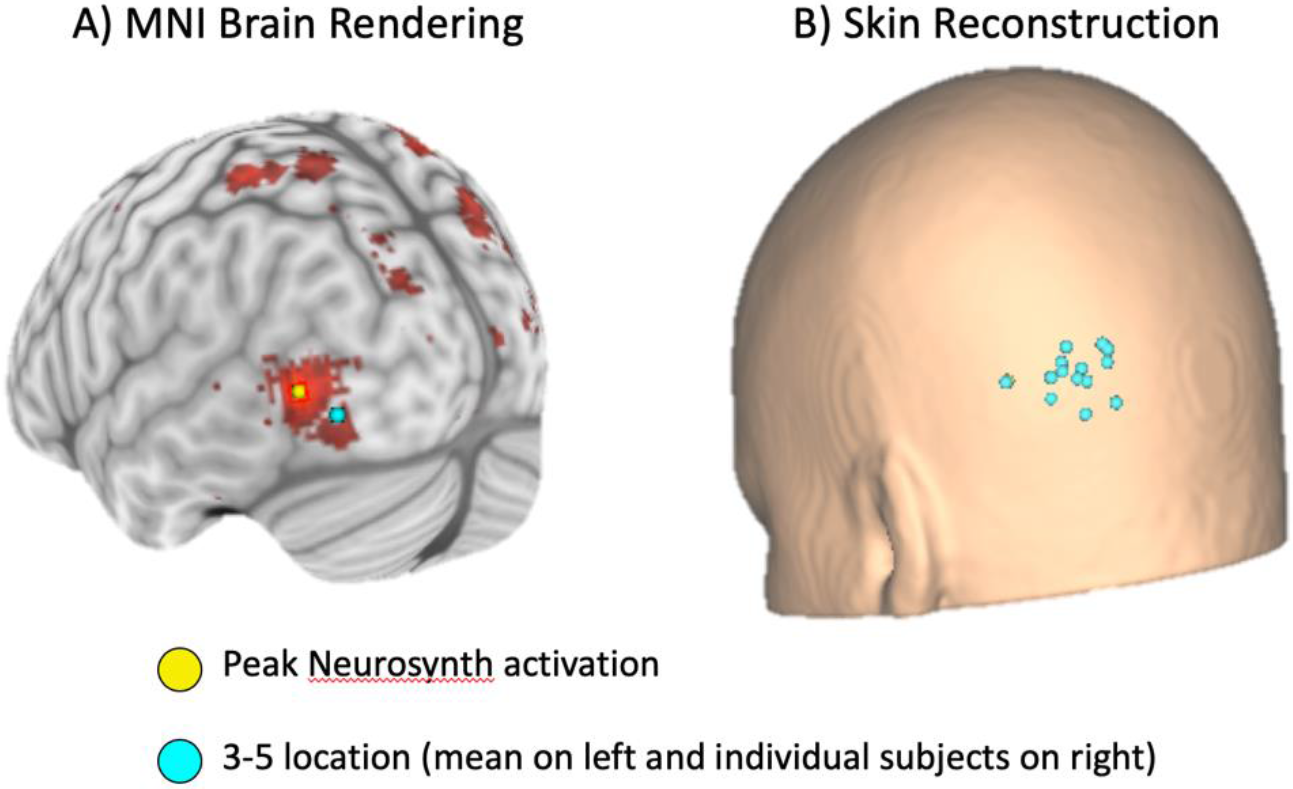
Location of stimulation sites. A) Neurosynth target (yellow dot) on “motion” probabilistic map and mean 3-5 target (cyan) shown on an MNI brain rendering. B) Skin rendering with individual subject’s 3-5 locations.

## DISCUSSION

This study assessed the effects of an online paired-pulse TMS protocol delivered over the left hMT+ at two distinct scalp targets as participants performed a motion direction discrimination task. This stimulation was expected to impair task performance, reducing accuracy in motion direction discrimination. While this task showed reliable parametric behavioral effects with increasing accuracy at higher coherence levels, there were no effects of ppTMS applied using either targeting method. We discuss possible explanations for this null finding below.

The results of the current study are in contrast with previous studies reporting modulation of hMT+ after ppTMS with different parameters. Decreased accuracy in motion perception has been observed when ppTMS (ISI: 5 ms) in 15 participants was applied 42 and 10 ms prior to stimulus presentation [20] or in five subjects 60–80 ms after stimulus offset [21]. Fast and slow moving dots have been disrupted selectively by ppTMS (ISI: 26.7 ms) applied in 12 subjects approximately 50 ms and 80 ms after stimulus onset respectively [19]. Furthermore, the capacity of predicting motion signal was impaired when ppTMS (ISI: 40 ms) was applied in 17 subjects, 13 ms before stimulus presentation [22]. Finally, ppTMS (ISI: 100 ms) to 36 subjects, applied 100 ms after stimulus onset, disrupted interception timing during an visual interception task [23]. Given that the current sample size of 15 is typical of these studies, and that the motion task was individually calibrated to maximize psychometric sensitivity, it is notable that the current study was not able to achieve similar active versus sham effects, as reported previously.

Factors including task sensitivity to TMS [33] and/or TMS parameters could have contributed to the negative results. For instance, the duration of the visual stimulus presentation used in this experiment may have been suboptimal to support ppTMS aftereffects. In particular, at 150 ms, the total duration of the motion stimulus presentation is somewhat longer than for other designs that have used single pulse TMS to disrupt motion perception [21, 30, 34]. In many of those studies, durations shorter than 100 ms were used under the logic that longer durations of motion stimuli may outlast the effective duration of modulation induced by TMS. This logic is further accentuated by Beckers and Zeki (1995) who highlighted that there are two paths to process motion perception, a fast pathway with direct access hMT+ and a slow one that arrives first to the striate cortex (V1), delivering information to hMT+ around 50 ms later [30]. It is possible that if there was an effect due to the ppTMS stimulation in this study, it may have been compensated for by the delayed action of the slow pathway, such that no difference was seen between active and sham stimulation.

In a similar vein, the timing of delivery of the TMS pulses with respect to the stimulus onset may have influenced the results. According to the literature, the critical time window to disrupt motion perception with TMS applied over the hMT+ extends from −40 to 200 ms around the stimulus onset [20, 21, 30, 34]. While there is considerable heterogeneity in the experimental parameters used in these studies several have shown that disruption in motion perception can be achieved with single pulses in an early period ranging from −40 to 0 ms before stimulus onset [20, 34, 35]. In the current protocol, paired-puse stimulation was delivered in the interval from −150 to −50 ms before the onset of the motion stimulus under the logic that persistent inhibitory effects of this TMS intervention will last for several hundreds of milliseconds and therefore overlap with the task duration. However, late cortical disinhibition, which peaks approximately 200 ms after a suprathreshold pulse [36], may have interfered with the intended inhibitory effects of the pulses, yielding a mixture of inhibitory and facilitatory effects. Thus, this long-interval ppTMS paradigm may have been suboptimal for inducing robust modulatory effects in motion perception.

An important consideration in any TMS study is the spatial positioning of the coil relative to the desired cortical target. With regard to motion-sensitive cortex in humans, several approaches can be adopted. These range from scalp measurement, to meta-analytic group coordinates (the so-called “probabilistic approach”), to individualized targets based on anatomical or functional neuroimaging. A commonly used scalp coordinate-based approach is the 3-5 cm rule [27, 28, 30], in which the coil is placed 3 cm dorsal and 5 cm lateral to the inion. This was the first targeting approach used in the current study. However, it is widely appreciated that such targeting does not scale with the head size of the participant, and therefore a second approach was implemented in which targeting was based on the peak activation obtained from the NeuroSynth meta-analytic database. In this case, the participant’s head was co-registered with a default scalp rendering from the MNI atlas and stereotactic navigation was used to guide coil placement. As such, this approach scales with the head size of the participant. While neither of these targeting approaches led to active-versus-sham behavioral effects, it is interesting that they did produce markedly different coil placements across individuals. In particular, these two approaches were offset from each other by an average of 2.9 cm, with an overall spread of several centimeters. Nonetheless, future studies may wish to capture individual variability in the functional location of hMT+ using fMRI-based targeting, as it has been shown that the improved precision can lead to stronger effect sizes [37].

Another final parameter choice that may have impacted the effects in this study was the decision to scale stimulation intensity at 120% of resting motor threshold. While such suprathreshold stimulation is common in TMS, it is based on sensitivity that is not derived specifically from the visual cortex and may have undermined potential behavioral results [38], suggesting that TMS motor thresholds cannot be assumed to be a reliable guide to the excitability of visual cortex. This suggests that more systematic investigations of the reliability of trial-wise TMS in eliciting consistent visual percepts with varying targeting approaches is deeply necessary. Nonetheless, other choices, such as stimulating at intensities that produce visual phosphenes, may have led to confounding perceptual effects wherein the phosphene may have interfered with motion sensitivity.

In conclusion, the present study highlights the complexity of TMS effects on visual perception and underscores the need for further dose-response studies in TMS literature to better understand the underlying neural mechanisms. This study can therefore serve as a reference point for future studies that systematically vary stimulation parameters to explore how these parameters modulate TMS effects.

## Notes

**Funding:** Research reported in this publication was supported by the National Institute of Mental Health of the National Institutes of Health under Brain Initiative Award Number RF1MH114253. The content is solely the responsibility of the authors and does not necessarily represent the official views of the National Institutes of Health.

